# Multiscale Quantification of Hemispheric Asymmetry in Cortical Maps Using Geometric Eigenmodes

**DOI:** 10.1101/2024.10.31.621232

**Authors:** Alicia Milloz, Jacob Vogel, Anders Olsen, James C Pang, Olof Strandberg, Toomas Erik Anijärv, Erik Stomrud, Sebastian Palmqvist, Nicola Spotorno, Rik Ossenkoppele, Dimitri Van De Ville, Oskar Hansson, Hamid Behjat

**Author notes:** Shared last-authorship/correspondence; e-mail: {, }.

## Abstract

Hemispheric asymmetry is a universal property of brain organization with wide implications into brain function and structure, and diseases. This study presents a laterality index for characterizing hemispheric asymmetries that underlie cortical maps using geometric eigenmodes derived from human cortical surfaces.We develop a generalized design to quantify asymmetries across various cortical spatial scales. While the design is individual-specific, we implement normalization steps to enable unbiased comparisons across individuals. As a proof of concept, we validated the method on cortical maps of 545 subjects across two datasets, using fMRI maps of healthy individuals and tau-PET maps of patients across the Alzheimer’s disease continuum. Our results reveal that cortical regions in different canonical functional networks have connectivity patterns that entail different degrees of hemispheric asymmetry. Moreover, aggregates of the pathological tau protein manifest subtle asymmetries at varying spatial scales along the disease continuum.

## 1. INTRODUCTION

Hemispheric asymmetry is a fundamental characteristic of brain organization and function, manifesting in various forms and scales, with a notable example being the structural and functional lateralization of language function [1]. Whilst studies have shown that hemispheric asymmetry is a feature of healthy brain structure and function, it has been found that brain asymmetries could also provide unique insights into various brain disorders such as Alzheimer’s disease [2], [3]. However, traditional methods for measuring hemispheric asymmetry rely on aggregate measures of brain morphology, like surface area and grey matter volume [4], which are only sensitive to single spatial scales. A growing body of research has suggested the potential of employing brain eigenmodes as a powerful, generalized framework to study the multiscale organization of the structure and function of the cortex [5]–[9]. The beauty of eigenmodes is that they provide a generating basis that spans the full range of spatial scales that can be manifested in empirical brain maps. In fact, multiscale asymmetries in the eigenmodes of brain geometry (aka geometric eigenmodes) have been shown to capture the individual-specific properties of healthy brains, akin to a fingerprint [10], and the links between brain structure and clinical symptoms in a cohort with early psychosis [11].

In this paper, we extend previous eigenmode-based approaches to develop a novel method that can characterize hemispheric asymmetries underlying brain cortical maps across multiple scales. In particular, we introduce a subject-specific, multiscale laterality index to quantify hemispheric asymmetry in cortical maps. We validate the method on cortical maps across 545 subjects from two datasets: (i) functional MRI seed-based connectivity maps in healthy individuals; and (ii) PET maps that measure aggregations of pathological tau protein in patients across the Alzheimer’s disease continuum.

## 2. MATERIALS AND METHODS

Fig. 1 shows an overview of the proposed method for multiscale quantification of laterality in cortical maps using subject-specific geometric eigenmodes. In the following, we describe the datasets used in this work, provide a brief outline of how geometric eigenmodes are extracted and used to decompose cortical maps, and finally present the proposed method for quantifying lateralization in maps.

**Fig. 1.**
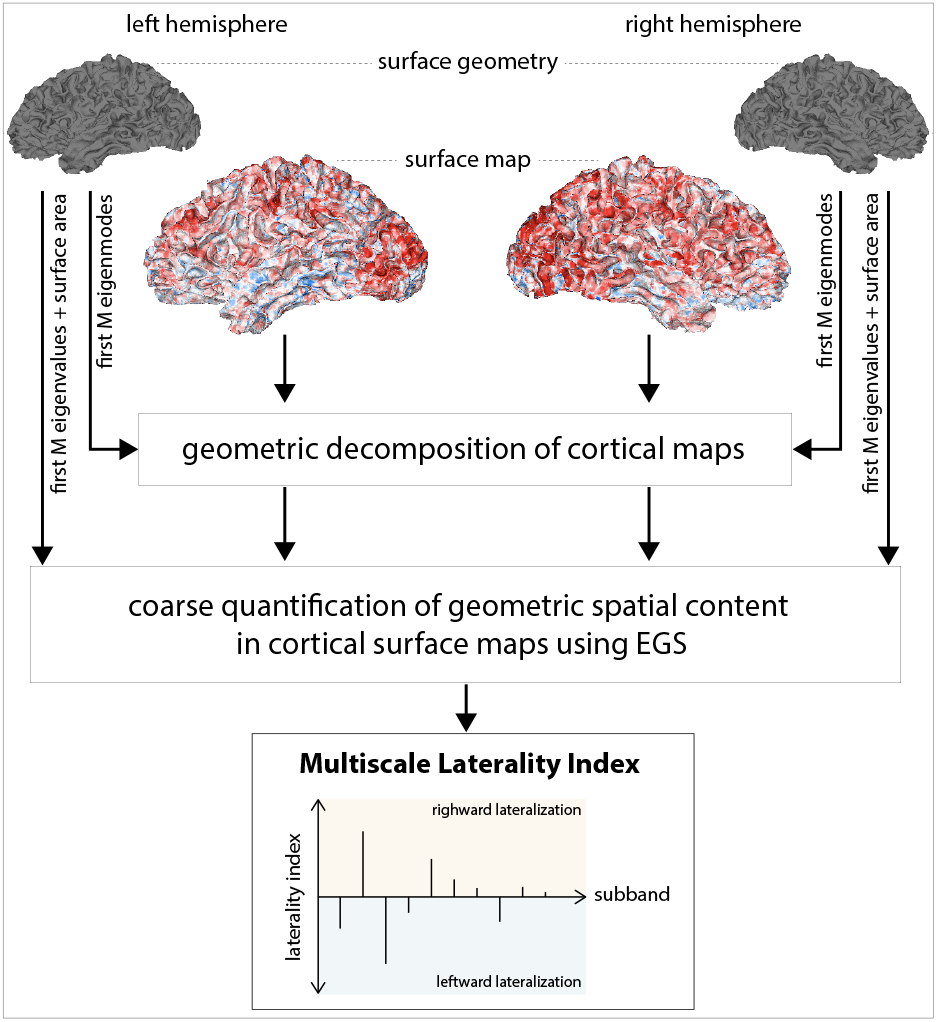
Overview of the proposed method for deriving a multiscale laterality index for any given cortical map.

### 2.1. Dataset

#### 2.1.1. Human Connectome Project

We used the minimally preprocessed structural and resting-state functional MRI data from the Human Connectome Project [12], [13], specifically, the 100 Unrelated Subjects set. As a proxy of seed-based functional connectivity, we constructed stationary co-activation pattern (CAP) map associated with cortical regions defined by the Schaefer atlas [14], classified into seven canonical functional networks [15], as in [16]. Here we focused only on the right hemisphere, consisting of 200 cortical regions; two subjects were excluded from our analysis due to errors in parcellating the cortices. Volumetric CAPs constructed in subject-space were projected onto the cortical surface of each subject—surface meshes with on average 139,137 *±* 12,551 vertices.

#### 2.1.2. BioFINDER-2

We used data from the Swedish BioFINDER-2 Study^1^ cohort [17], a prospective cohort study that aims to identify biomarkers for neurodegenerative diseases. We studied 447 individuals aged over 50, having a structural T1 MRI scan and an [^18^F]RO948 PET scan [18] that indicates tau protein content (quantified as Standardized Uptake Value Ratios [SUVR] maps). Subjects were classified [19] by cognitive status (cognitively unimpaired [CU], mild cognitive impairment [MCI], and Alzheimer’s disease [AD]), and biomarker status (amyloid-*β* positive [A+] or negative [A-], based on CSF test [20]; tau positive [T+] or negative [T-], based on a temporal meta ROI threshold of 1.36 [21]). We classified the subjects into four groups denoted as: `CU/A-/T-` (N=88), `CU/A+/T-` (N=94), `MCI/A+/T+` (N=93), and `AD/A+/T+` (N=172).

### 2.2. Geometric decomposition of spatial maps

Given a surface mesh representation of a cortical hemisphere— generated from a T1-weighted MRI image via e.g. the FreeSurfer software suite [22]—one can construct its corresponding Laplace-Beltrami Operator (LBO) Δ—which captures local spatial information and curvature of the cortical surface—and eigendecomposed as [5] Δ**Ψ** = *−***ΛΨ**, resulting in a set of geometric eigenmodes

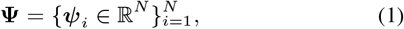

and an associated set of eigenvalues

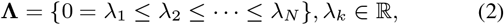

which defines the geometric spectrum of the hemisphere; *N* is the number of vertices in the mesh, unique to each hemisphere. The eigenmodes are ordered in increasing spatial frequency (decreasing spatial wavelength), each entailing a neuroanatomical cortical spatial pattern at a specific scale defined by its eigenvalue (cf. Fig. 2); in the limit of vanishing cortical foldings, geometric modes approach spherical harmonics, linking each geometric eigenvalue to a physical spatial scale dependent on the total surface area of the mesh (i.e., radius of sphere) [5], [23]. We used the LaPy Python package [24] implementation, and only computed the leading *M* eigenmodes, where *M* ≪ *N* . A spatial map defined on the cortex **y** ∈ ℝ^*N*^ can be decomposed using a general linear model (GLM) as:

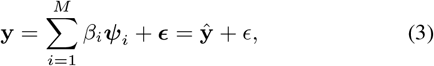

where *β*_*i*_ is the coefficient quantifying the contribution of *Ψ*_*i*_ to the map and *ϵ* is a constant vector representing the error. To prevent any bias on estimated weights when comparing across hemispheres and subjects, all spatial maps are l2-normalized before decomposition.

**Fig. 2.**
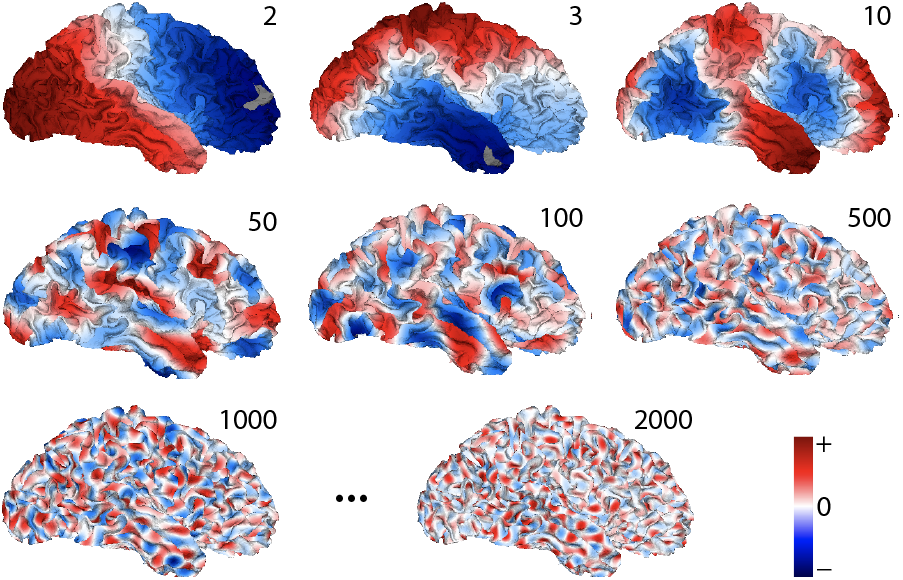
Geometric eigenmodes of a representative subject’s right hemisphere; *i* = 2, 3, …, 2000 indicate eigenmode index *Ψ*_*i*_.

### 2.3. Multiscale laterality index

Our aim is to develop a laterality measure to quantify hemispheric asymmetries in cortical spatial maps. The basis of this measure is the estimated 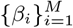 from the map on the left and right hemispheres. Importantly, we want this measure to be normalized such that it can be compared across subjects. There are two complications to this goal. (1) Differences in surface areas result in a multiplicative scaling of the entire eigenspectrum, which may be directly accounted for by applying a rescaling function [24]. (2) Each geometric spectrum **Λ** is unique in size (*N*) and eigenvalue distribution (λ_*i*_). This variability creates a potential bias if one aims to make one-to-one comparisons between *β*_*i*_ not only across subjects but also between the left and right hemispheres of the same subject. To overcome this challenge, inspired by prior work [25], we propose working at the level of spectral sub-bands [26] of the geometric spectrum—obviating the need for one-to-one comparisons between *β*_*i*_—resulting in a measure that is normalized for comparison across subjects [27]. Let 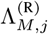 denote the sub-spectrum of the right hemisphere of subject *j* consisting of the first *M* eigenvalues, 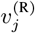 the surface area of the hemisphere, and 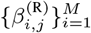 the GLM coefficients associated to a spatial map defined on the hemisphere. Let 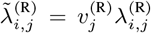 denote the surface-area-corrected versions of 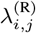 [24], a correction that removes the bias related to differences in hemisphere size [28]. Similarly, we define the aforementioned variables for the left hemisphere, replacing ·^(R)^ with ·^(L)^ in the notations.

#### Definition (Ensemble Geometric Sub-bands [EGS])

Given the surface mesh representations of the left and right hemispheres of *J* subjects, a set of Ensemble Geometric Sub-bands associated with the first *M* eigenmodes can be defined via a system of spectral kernels as

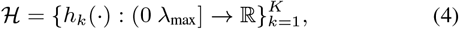

where *K* is the number of sub-bands, and the maximum spectral value λ_max_ is defined as

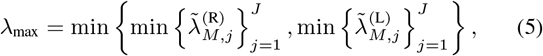

see Fig. 3(a). A special case of EGS is obtained when sub-bands have a flat weight and are non-overlapping, that is:

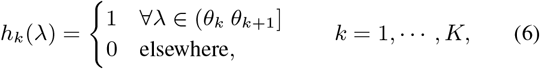

where 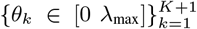 denote spectral positions that satisfy *θ*_1_ = 0, *θ*_*k*_ < *θ*_*k*+1_, and *θ*_*K*+1_ = λ_max_; specifically, each spectral pair 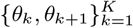 specifies the lower and upper bounds of a subband; see Fig. 3(b).

**Fig. 3.**
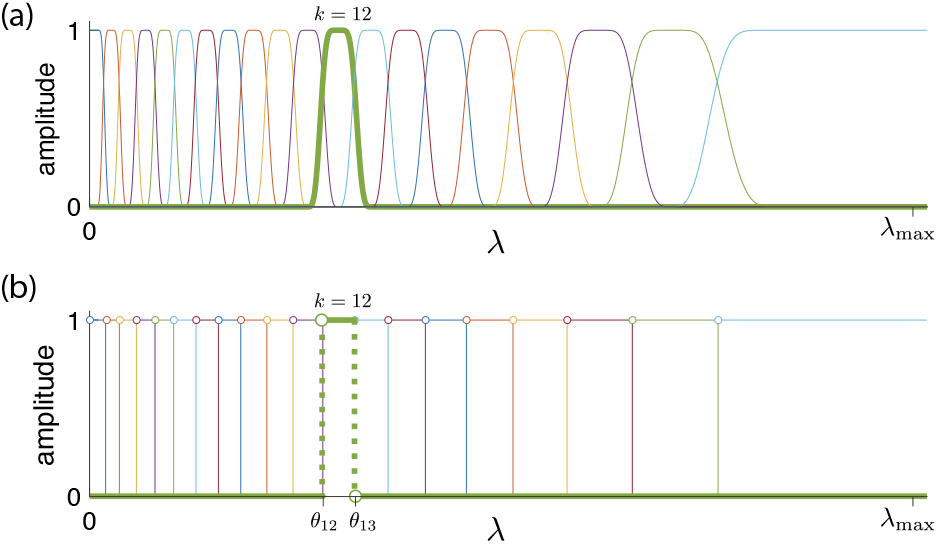
Ensemble geometric sub-bands (EGS). (a) An example system of EGS; cf. Eq. (4). (b) A special-case system of EGS with non-overlapping sub-bands; cf. Eq. (6). There are 20 sub-bands in both (a) and (b), with sub-band 12 highlighted for comparison.

For each dataset (fMRI CAP maps and tau-PET maps), we determined the number of eigenmodes *M* to reach an overall reconstruction accuracy (average across maps and subjects) of 70% or higher, exceeding that used in prior work [5], [8], [9]; reconstruction accuracy was determined by the Pearson correlation coefficient between **y** and **ŷ** in (3). We found the approximate suitable *M* to be 2000 for fMRI CAPs and 500 for tau-PET maps, which, intuitively, reflects that the latter maps entail smoother spatial profiles than the former. We also designed an unique EGS for each dataset. This was done using a signal-adapted design strategy [29], which defines sub-bands to capture an equal average amount of signal energy. Since our objective is to quantify the spatial profile differences between the left and right hemispheres, we considered the ensemble difference between the energy content of the left and right hemisphere maps as the measure to equalize across sub-bands. Fig. 3(a) shows the designed system of kernels for fMRI CAPs based on [29], which we then simplified to the non-overlapping version shown in Fig. 3(b); EGS for tau-PET maps (not shown) entailed a similar spectral partitioning to that of fMRI CAPs but had notably narrower sub-bands at the lower end of the spectrum.

#### Definition (Multiscale Laterality Index [MLI])

Given the left and right hemisphere surface representation of a subject’s cortex, and a cortical spatial map defined on each surface, the laterality in the map can be quantified by a multiscale laterality index as

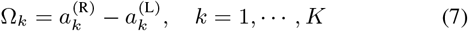

where 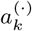 is defined as

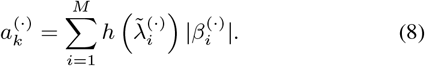

### 2.4. Permutation testing to determine significance of MLI

We used permutation testing to determine the significance of the MLI. For each map, we generated 1000 surrogate maps by randomly shuffling the elements in the map. The MLI of each surrogate map was then determined, resulting in a null distribution of MLI for each sub-band. A *p*-value was then determined for each sub-band at the significance level of *α* = 0.01*/K*, where the denominator indicates Bonferroni correction of multiple comparisons across sub-bands.

## 3. RESULTS AND DISCUSSION

Fig. 4 shows the results on fMRI CAPs. Fig. 4(a) shows that a large proportion of CAPs entail a statistically significant laterality index (cf. Section 2.4). The proportion not only varies across networks and sub-bands but also across subjects, suggesting subject-specific lateralization idiosyncrasies in functional connectivity patterns. Fig. 4(b) illustrates the degree of asymmetry in CAPs, revealing a landscape of functional connectivity that substantially varies across different cortical regions and networks. Frontal, posterior parietal, and lateral temporal cortices show the highest degree of asymmetry.

**Fig. 4.**
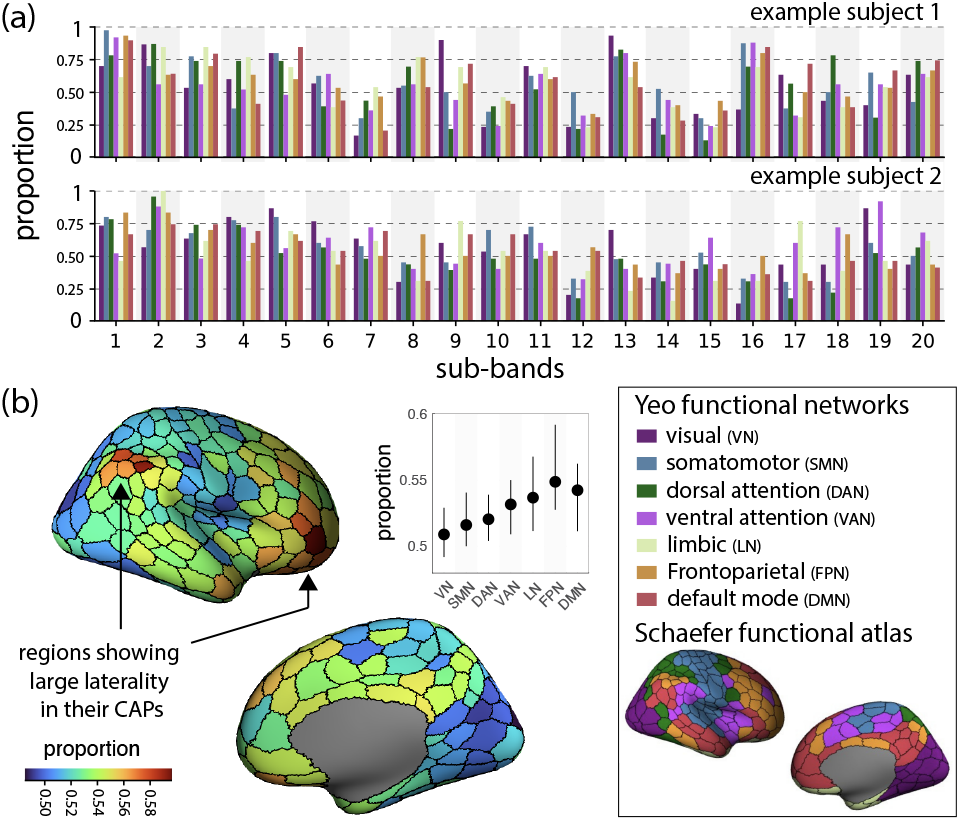
MLI of fMRI CAPs in healthy individuals. (a) Proportion of statistically significant MLI across networks for two example subjects. (b) Proportion of significant MLI across the cortex; proportions computed across subjects and sub-bands.

At the network level, these regions belong to the intrinsic frontoparietal and default mode networks. In contrast, cortical regions within the visual, dorsal attention, and somatomotor networks entail a lower degree of laterality. This suggests an increasing connectivity lateralization along the primary gradient of brain organization, from low-level (somatomotor, visual) to higher-level association cortices. Future research should explore the functional implications of these asymmetries and their role in human cognition and behavior.

The analysis of tau asymmetry within the Alzheimer’s Disease (AD) continuum reveals insights into the differing patterns of tau asymmetry across clinical and cognitively unimpaired groups. Fig. 5(a) shows that patients with AD and MCI are associated with tau asymmetry on wider spatial scales (up to sub-band 10). This suggests that the detected tau asymmetry may be more widely distributed in the brain for these groups. In contrast, asymmetry in cognitively unimpaired individuals with lower tau burden predominantly appears at finer spatial scales (sub-bands 13 to 20), suggesting that the observed asymmetries may differ in nature, potentially reflecting off-target tracer binding or non-pathological imaging features. Fig. 5(b) shows the amplitude of MLIs across sub-bands and groups, highlighting also the variability across subjects. Group `CU/A-/T-` shows significant leftward asymmetry in sub-bands 5, 6 and 17, whereas group `CU/A+/T-` shows significant rightward asymmetry in sub-bands 14 and 16. The MCI and AD groups both show significant leftward asymmetry in sub-band 4, whereas the AD group also shows significant leftward asymmetry in sub-band 5. Overall, laterality indices are larger in clinical groups (MCI and AD) than in cognitively unimpaired groups. This indicates that lateralization magnitude may be a better indicator of cognitive status than its direction. Additionally, this may reflect selection bias in the cohort, as individuals with more lateralized pathology are more likely to have overt symptoms, leading to memory clinic referrals and MCI/AD diagnoses. Fig. 5(c) shows the relationship of laterality index of sub-band 4 with age and mean tau PET (mean tau PET SUVR value across a meta ROI including Braak stages 1-4 [30]). We found that a greater leftward tau asymmetry is associated with a higher mean tau SUVR; linear regression analysis coefficient is -1.34, and p <0.05. We observed no significant linear association between the laterality index and age.

**Fig. 5.**
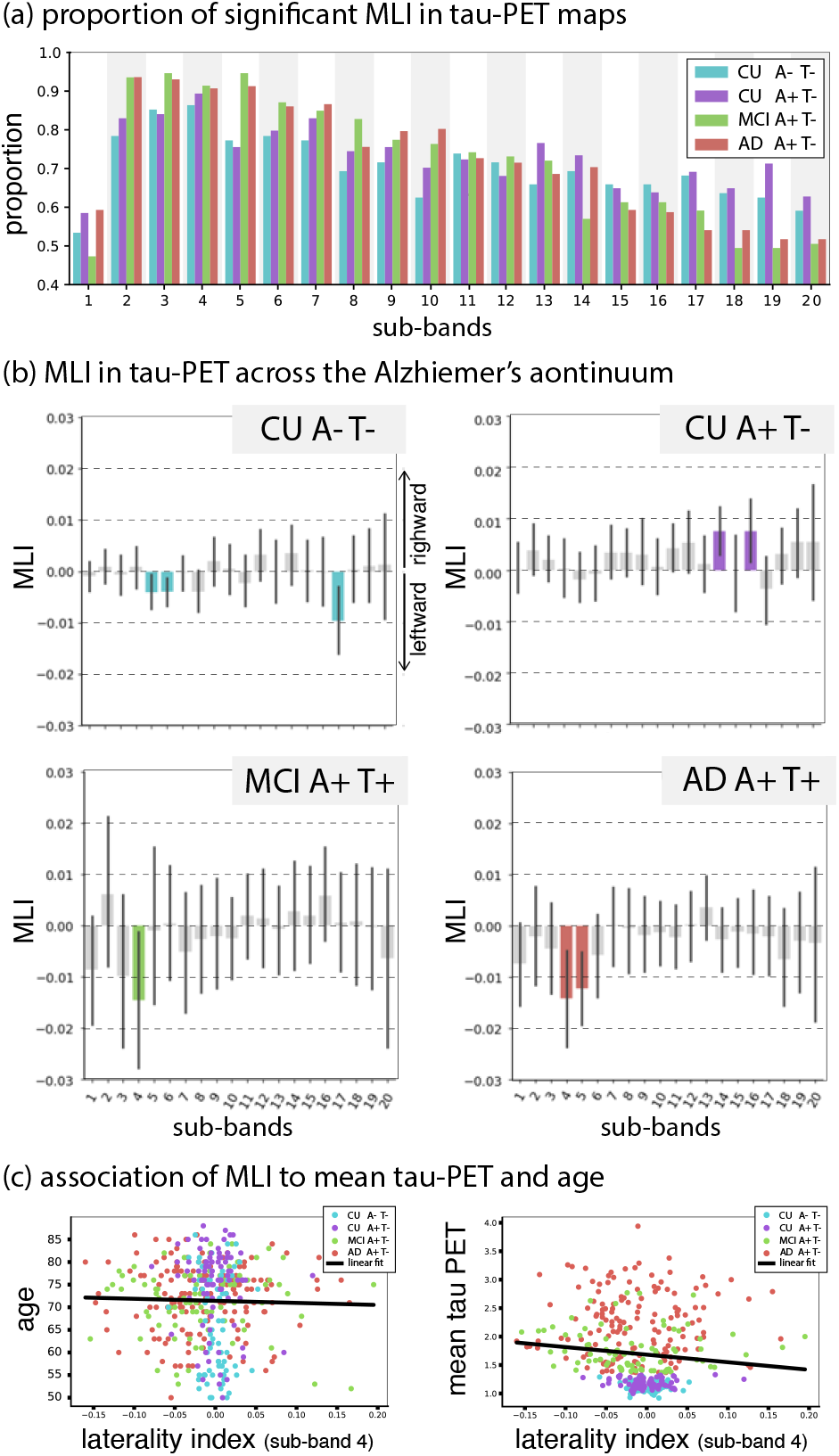
MLI of tau PET along the Alzheimer’s disease continuum. (a) Proportion of significant laterality indices across sub-bands and groups. (b) Amplitude of laterality indices, where each bar reflects the mean and the whisker represents the confidence interval; a subband is marked with color if its laterality index amplitude is significant — i.e., confidence interval does not cross zero (not corrected for multiple comparisons) and is marked with gray otherwise. (c) Relationship between laterality index of sub-band 4 with age (left) and mean tau PET SUVR (right), respectively.

Our results suggest the potential of using a multiscale approach in quantifying asymmetry in brain maps, which can capture a wide range of spatial variations across spectral scales. Here, we used permutation tests to determine the significance of MLI, but future work may consider integrating suitable statistical tests in the definition of the index. Moreover, the number of sub-bands and the strategy used to define EGS are design parameters that require further investigation to obtain optimal sub-bands. In particular, given the generalized formulation in the design of EGS, different designs of multiscale systems of spectral kernels [31] can be explored in future work. Computationally, the proposed MLI can be extended to bypass explicit computation of individual eigenmodes, and thus *β*_*i*_, leveraging graph signal processing spectral filtering principles [32] to determine energy associated to a coarse set of spectral kernels that can be approximated as polynomials. Lastly, the proposed method can be extended to leverage other forms of brain eigenmodes, such as eigenmodes of high-resolution connectomes [33] or voxel-wise cortical [34], white matter [35], or whole-brain [6] graphs.

## 4. CONCLUSIONS

We proposed a novel method to quantify asymmetry in any cortical map at multiple spatial scales using geometric eigenmodes. The resulting index is subject-specific, relying on each individual’s cortical geometry, but is normalized for seamless comparison across subjects despite using individual-specific eigenmodes. By studying spatial patterns in cortical maps at the level of sub-bands rather than at the level of individual eigenmodes, we, firstly, overcome the challenge of matching eigenmodes across left and right hemispheres, and across subjects, and, secondly, obviate the need to study and interpret the role of separate eigenmodes. As a proof-of-concept, we used the method to reveal how fMRI co-activation patterns entail different degrees of hemispheric asymmetry across the cortex and functional networks, and how this varies across geometric sub-bands. Furthermore, the application of the method to tau-PET maps highlights a general leftward asymmetry in MCI and AD, with a notable substantial magnitude at a specific spatial scale. The proposed method opens a rich avenue for future research into the interplay between brain structure and its manifested function and pathology.

## 5. COMPLIANCE WITH ETHICAL STANDARDS

This research study was conducted retrospectively using human neuroimaging data ethical approval for acquisition of which have been previously obtained. The authors certify that they have no conflict of interest to report in regards to the subject matter discussed in this paper.

## Footnotes

1 https://biofinder.se

